# Harnessing the Potential of Cape Wild Edible Plants: Nutritional insights, Gaps and Priorities

**DOI:** 10.1101/2025.07.23.666301

**Authors:** Nicola Kühn, Loubie Rusch, Michelle Rorich, Frances Storey, Emma van der Meulen, Laura Pereira

## Abstract

Rationale: Wild edible plants of the Greater Cape Floristic Region (GCFR) of South Africa have piqued recent interest from multiple sectors due to their potential to contribute to the Western Cape’s food systems. However, cultivation efforts remain hindered due to a lack of nutritional knowledge.Methods: This systematic literature review establishes nutritional knowledge gaps and priorities for a selection of the region’s wild edible plants and uses the existing nutritional data to summarize the proportionate contributions that these species can make to daily dietary requirements.Key results: We find that winter rainfall/GCFR species are poorly researched, and more so than summer rainfall species in terms of a) nutritional status studied and b) number of key nutrients tested in each species. We also identify key winter rainfall species (e.g. *Carpobrotus edulis, Tulbaghia violaceae and Trachyandra falcata*) that show high potential to address nutritional deficits of the region.Conclusion: Although we find nutritional data to be sparse, that which exists suggests high nutritive potential of wild edible plants of the GCFR. We also identify priority species for future nutritional study and cultivation to enhance local food systems using the bounty offered by this biodiversity hotspot.

**Societal impact statement:** This study highlights the potential of wild edible plants in the Greater Cape Floristic Region (GCFR) to enhance food security and nutritional health in the Western Cape. By identifying nutritious species like *Carpobrotus edulis*, and poorly researched yet nutritious species like *Tulbaghia violaceae*, and *Trachyandra falcata*, we emphasize the role of GCFR edible plants in addressing deficiencies and promoting diversity of diets. Cultivating these plants can stimulate local agriculture, support sustainable practices, and empower communities. Ultimately, our findings advocate for reintegrating wild edible plants into local food systems, fostering food sovereignty and resilience while reducing reliance on global supply chains.

## 1. Introduction

Wild edible plants offer immense potential to strengthen global food systems, making them more nutritionally secure, climate-resilient, sustainable, and healthier. In areas where crop failures or food shortages occur, local wild edible plants serve as vital resources. At the same time, increasing the diversity of plants in diets is consistently associated with improved health outcomes across different regions and contexts (Gold & McBurney, 2012; Motti, 2022). Additionally, species adapted to local climates and soils are more likely to withstand climatic shocks and environmental changes, further enhancing their value in building resilient food systems. The adoption of wild edible plants also has additional societal benefits by empowering local market stakeholders, and connecting consumers with producers, thereby reducing reliance on global value chains and enhancing food sovereignty (Borelli et al., 2020). This makes the preservation and cultivation of wild edible plants, particularly in biodiverse regions which are shown to coincide with high diversity in edible plants (Pironon et al., 2024), even more critical.

The Greater Cape Floristic region (GCFR) in South Africa is the smallest yet most biodiverse hotspot on the planet. It supports approximately 11’423 species (Snijman, 2013) of which many are only found in this region. Around 1’740 of these species are edible plants yet very few are regularly consumed or cultivated (Welcome & Van Wyk, 2019). Of concern is that approximately 25% of the total species from the GCFR are threatened in the wild (Rouget et al., 2014) due primarily to extensive habitat degradation and loss from urban and large-scale agricultural expansion. Invasive species encroachment is also a key threat, while climate change is identified as a significant future threat (DEA (Department of Environmental Affairs), 2013; Ziervogel et al., 2014). Hence, worries arise that these combined threats might diminish the cultural and practical worth of the region’s edible plant diversity—a significant loss given their historical and potential future role in supporting livelihoods (Akinola et al., 2020).

For millennia, the local inhabitants of the GCFR relied on foraging of the region’s edible plants (Botha et al., 2020; Wehmeyer, 1986). The archaeological records even indicate that humans evolved to become modern based on their micronutrient and carbohydrate-rich diets drawn from the local landscapes (Singels et al., 2016, Singels et al., 2020). Post-colonially, local indigenous edibles of the region have become transposed by foods belonging to incoming food cultures, bringing with them the large-scale agricultural practices used to access them. Indigenous foods have become absent from the region’s local foodways, supply chains and knowledge systems despite their ecological adaptedness, cultural relevance, deep historical ties with humanity and potential nutritional value (Uusiku et al., 2010). This absence has been less apparent for the indigenous edibles in the rest of South Africa, in the summer rainfall regions, where edible plants have been cultivated on subsistence scales and thus remained key food sources (Akinola et al., 2020).

Today, the socio-economic circumstances of the majority living in the Western Cape is resulting in childhood stunting among 21.4 % of children under 5 years old [2017 data (UNICEF/WHO/World Bank, 2022)] and non-communicable lifestyle diseases being the cause of death among about 25% of the population in the Western Cape (Solomons et al., 2019), including but not exclusive to those living in poverty. Furthermore, past baseline studies have indicated poor iron (Motadi et al., 2023) and zinc status and vitamin A deficiency (Faber M et al., 2011), as well as seasonal deficiency in Vitamin D (Middelkoop et al., 2022). This places a burden on the Province’s health system. A question arises, whether an opportunity exists to tap into the biodiversity of locally adaptive foods of the region to address nutrition deficits.

The recent resurgence of interest in foraging and making use of local wild edibles in the Western Cape, running parallel with international trends, has been supported through improved understanding of how to grow and use a selection of local edibles (Rusch, 2021, 2022). However, formalised cultivation efforts of GCFR species has remained slow. A key reason for this is a lack of knowledge and synthesis of the knowledge that does exist on the nutrition of these species. Setting strategic and informed priorities for cultivation has thus proven difficult so far. This lack of information not only limits the extent to which their uptake can be promoted for the purpose of improving health, it is also information that interested consumers as well as the public bodies controlling health and safety regulations require to be displayed on foods promoted or sold in formalised food systems.

Alongside identifying plants that could mitigate against nutrient deficiencies in the region, establishing a baseline understanding of nutritional status or data deficiency of wild edibles, will promote the wider use of these species (for example, uptake from professional cooks which will in turn support small scale farmers) which will have knock on economic benefits to society.

Our study contributes to these knowledge gaps by synthesising existing nutritional data, including an understanding of potential contributions to daily dietary requirements, on a selection of wild edibles growing in the Greater Cape Floristic Region of South Africa. Additionally, we aim to identify nutritional information gaps and, in so doing, establish priority species for cultivation and thus reintegration into the food and knowledge systems in the Western Cape. We therefore hypothesise the following:

1. Species indigenous to the winter rainfall/Greater Cape Floristic Region are currently significantly under-researched and under-published in terms of nutritional status (in terms of research effort and nutrients considered).
2. For those winter rainfall/GCFR species with existing data, nutritional potential is high.

This baseline study thus sets out to establish and present the current nutritional information status of a selection of 22 local wild edible plants growing in the Greater Cape Floristic Region. It therefore intends to serve as a baseline from which to expand future work in support of both traditional knowledge systems as well as the sciences being used to uphold the potential of GCFR local wild foods to make social and ecological impact.

## 2. Methods

### 2.1 Species list

The species used in this study are those presented in the two publications “Cape Wild Foods: A Growers Guide” (Rusch, 2021) and “Cape Wild foods: a Cooks Guide” (Rusch, 2022) and listed in **Figure 1**. The inclusion/exclusion criteria of species can be summarised as those species found in the GCFR that are suitable for growing at household and small-scale cultivation scales. Some are wild food plants not previously propagated or cultivated and until recently only occurring in the wild, while others are found propagated and for sale in nurseries and are grown in public as well as private gardens. Not all of the species are endemic to the GCFR, with some occurring naturally in the summer rainfall areas, but acclimated to growing in the GCFR.

**Figure 1a).**
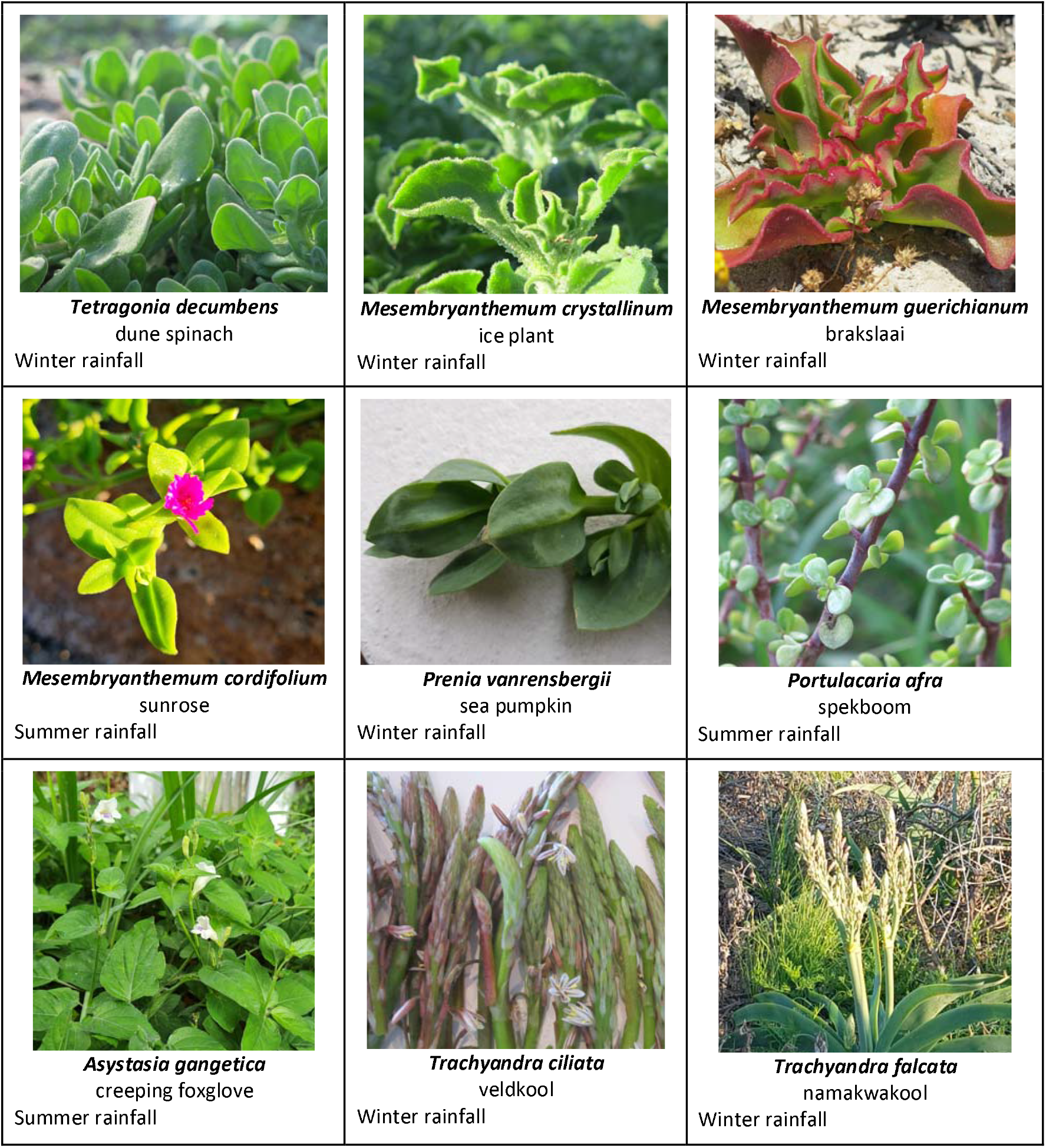
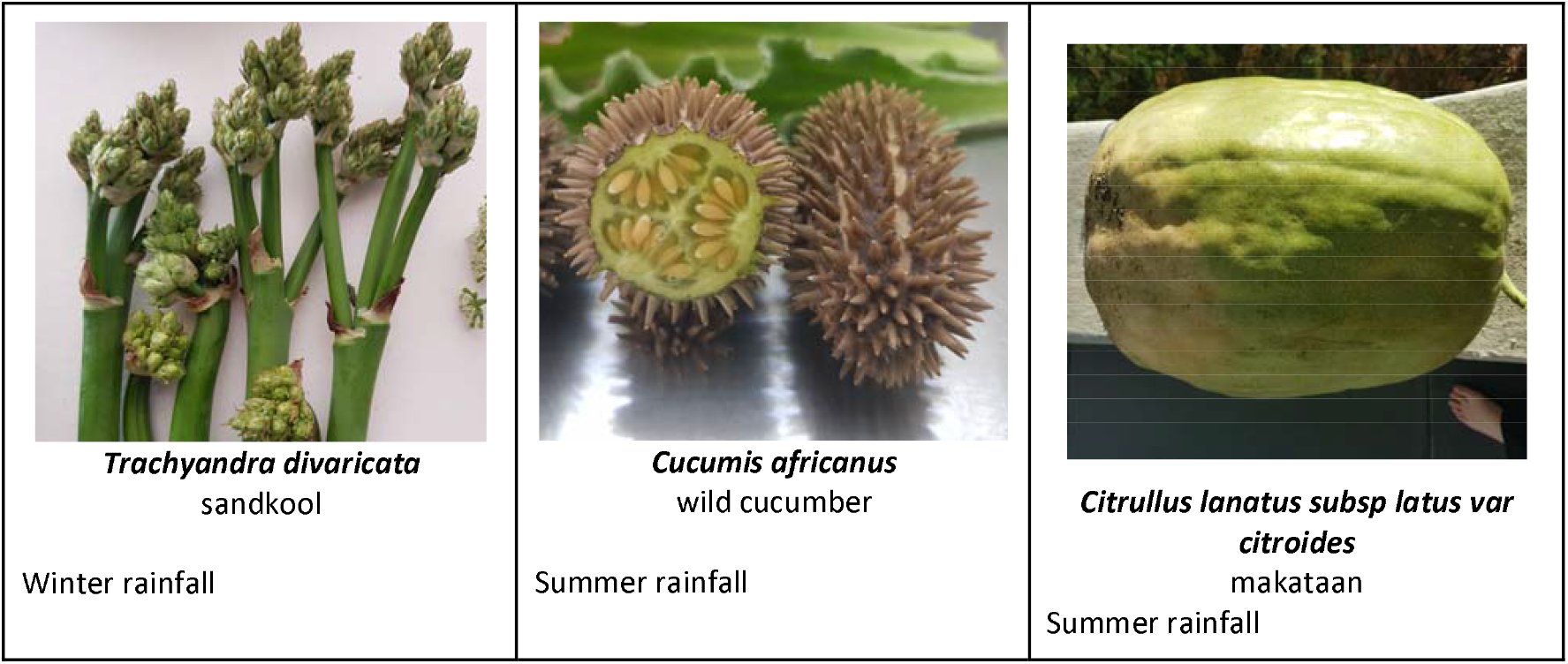
Leafy Greens, Succulents and Vegetables featured in this study (See **supplementary Table 1** for additional common names).

**Figure 1b).**
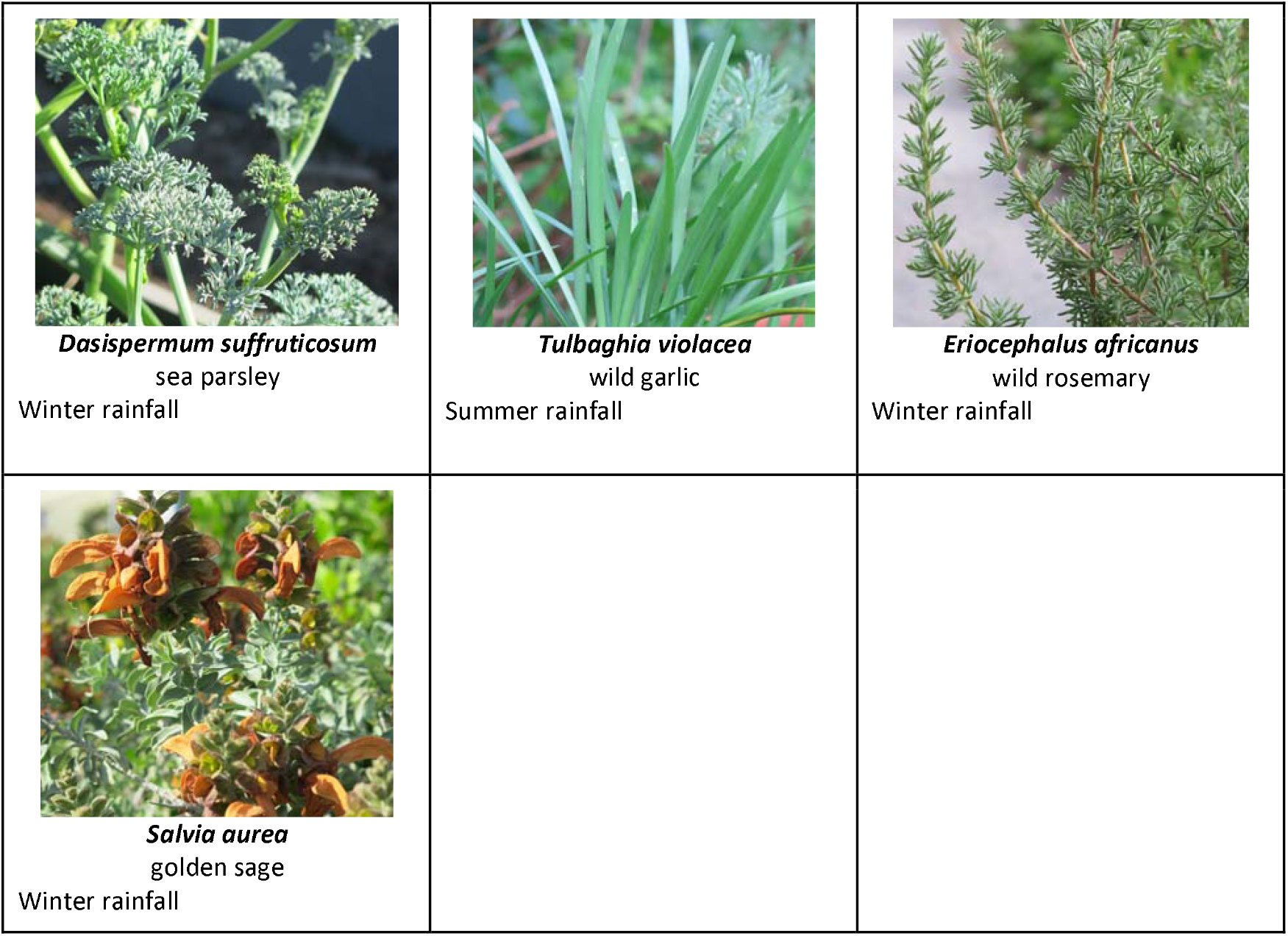
Culinary Wild Herbs featured in this study (See **supplementary Table 1** for additional common names).

**Figure 1c).**
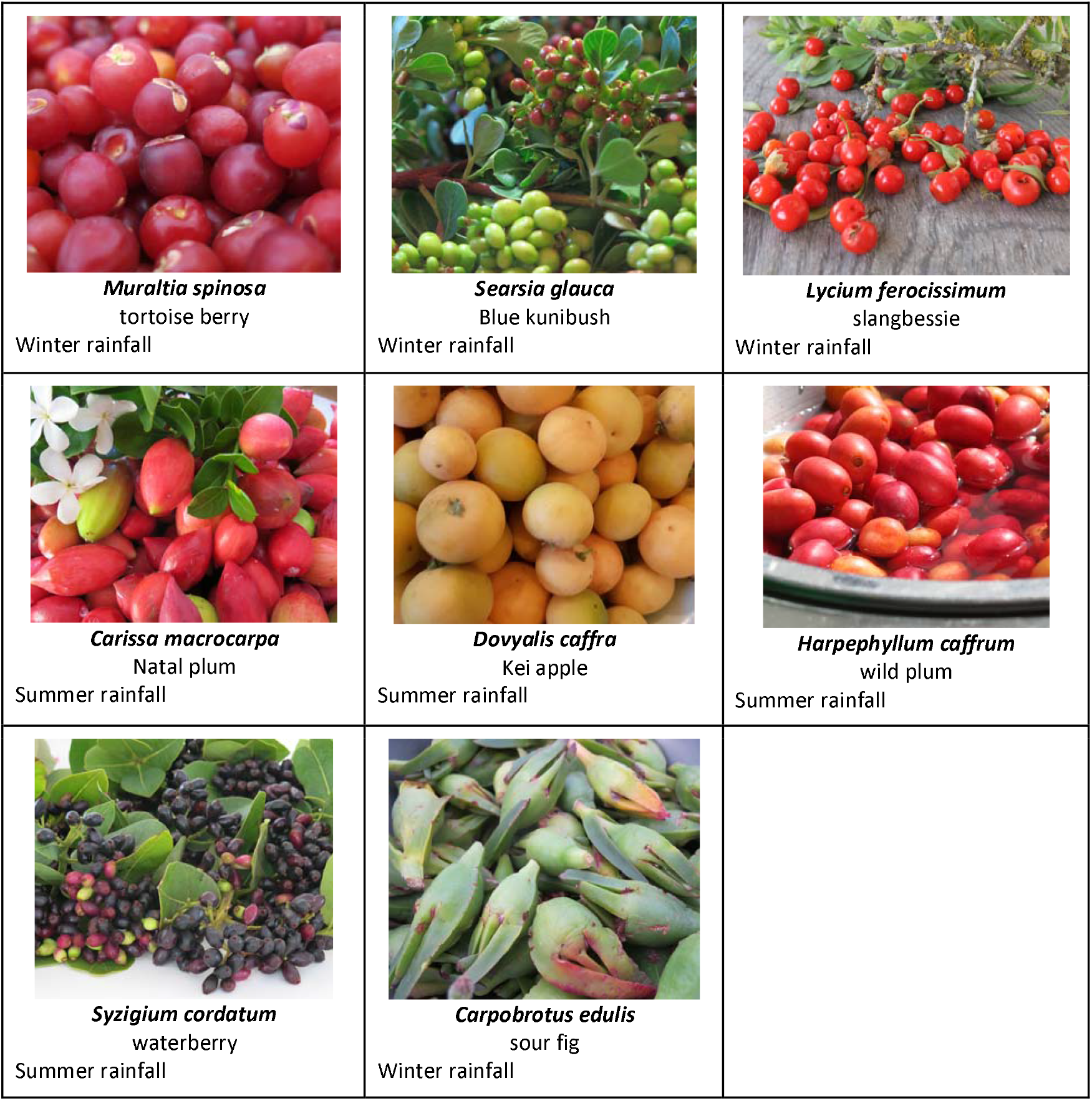
Soft Fruits and Drupes featured in this study (See **supplementary Table 1** for additional common names).

### 2.2 Literature Review

We conducted a literature review in the Google Scholar database to identify the most recent relevant advances in nutritional research (up until August 2022) on the selected species. We used the following combination of search terms: [the name of each species] + “∼nutrition”. The total number of search hits were 17’148 titles. We read the top 20 most relevant papers (as defined by the google scholar algorithm of relevance) for each species. If there were very high numbers of papers we focused on those between 2017-2022. If no papers were present, grey literature was also searched. From this reviewing process we filtered this list to 89 papers that met the criteria of containing nutritional information for our species, resulting in 133 records of nutritional status which were used to generate the database of nutrition status. We recorded the presence or absence of studies for each species. Upon identifying a study that presented data on nutritional status, we recorded this for the list of key nutrients (Macronutrients: Energy, Protein, Carbohydrates, Fibre, Fat, Sugar. Micronutrients: Calcium, Magnesium, Zinc, Iron, Vitamin A, Vitamin C, Vitamin D, Vitamin B9). If this first general search did not yield any information on these key nutrients, we expanded the search to [species name] + [key nutrient name].

#### 2.3.1 Analysis - Research Effort

From our literature review, we recorded the overall research effort including the total number of search hits per species and compared the winter vs summer rainfall origin species data. From this data we analysed the percentage of incidences of where the nutrient was studied proportional to the number of species of either winter or summer rainfall was considered so as to obtain a relative percentage of research effort for each nutrient by summer vs winter rainfall.

#### 2.3.2 Analysis - Key nutrients

To compare existing ranges of nutritional values of key nutrients of those species studied we conducted a comparative analysis among species. Those studies presenting data for a key nutrient of a species, were recorded alongside the unit it was measured in and the region it is indigenous to (GCFR/winter rainfall vs summer rainfall). For each record we then standardised each key nutrient value to have the same unit. This was used to calculate the percentage contribution of a nutrient relative to the daily recommended intake (defined for an adult female from the British Nutrition Foundation) of that nutrient, per portion of the species. The food portion used for this analysis for each species was determined by its category of being vegetable-like/succulent leaves/leafy greens (80g average portion), fruit/drupe/berry (80g average portion) or herbs/aromatics (10g average portion).

## 3. Results

### A: Nutritional research effort

Overall search hits suggest similarity between winter and summer species (average of 717 vs 710), however, only two species (*Carpobrotus edulis* and *Mesembryanthemum crystallinum*) account for the majority of these search hits for winter species, whereas summer species tend to have more evenly spread hits suggesting each species is more consistently studied (**Fig. 2a**). Species that lack any data on key nutrients for diets include *Mesembryanthemum cordifolium* for summer rainfall species and, *Prenia vanrensburgii, Trachyandra cilliata, Dasispermum suffruticosum, Searsia glauca, Mesembryanthemum guerichianum, Salvia aurea* and *Trachyandra divaricata* (**Fig. 2 b**) for winter rainfall species.

**Figure 2.**
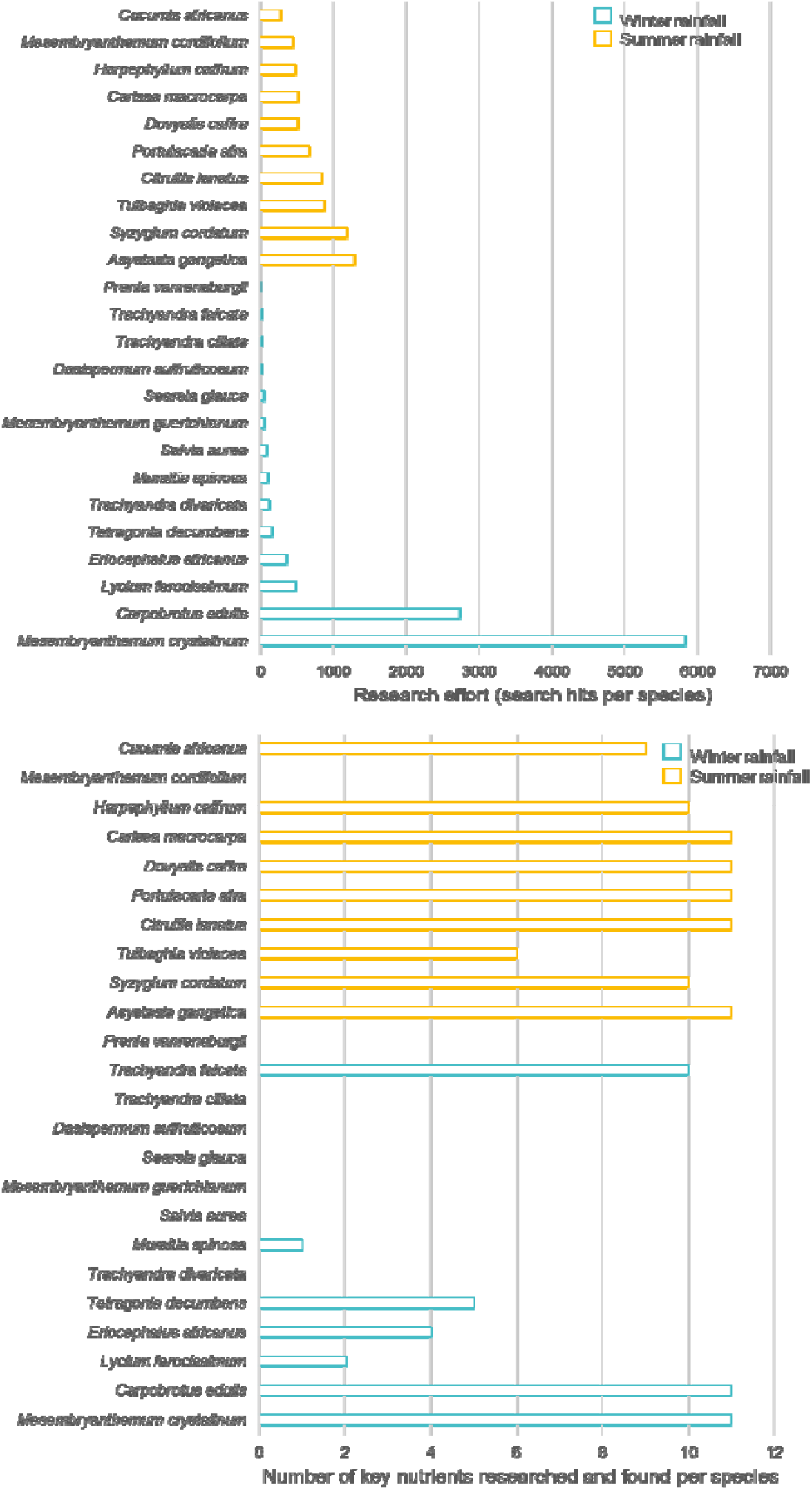
**a)** Research effort defined by the number of articles found (search hits) per species including the search terms of species name + ∼nutrition. **b)** The coverage of key nutrients in existing literature as defined by the number of key nutrients researched per species.

Research into nutrients (and some antinutritional components) indicate overall less relative research for winter rainfall vs summer rainfall species (**Figure 3**) (winter rainfall species studies make up one quarter that of summer rainfall species for energy, carbohydrate, fat, sugar, calcium and manganese and one half that of protein, sugar, fibre, magnesium, zinc, iron, Vitamin C, copper, chromium and phytochemicals). Notably, in both regions Iodine and Vitamins D, B9, E and K are under researched, however zero relevant data exists for winter rainfall species for these nutrients.

**Figure 3.**
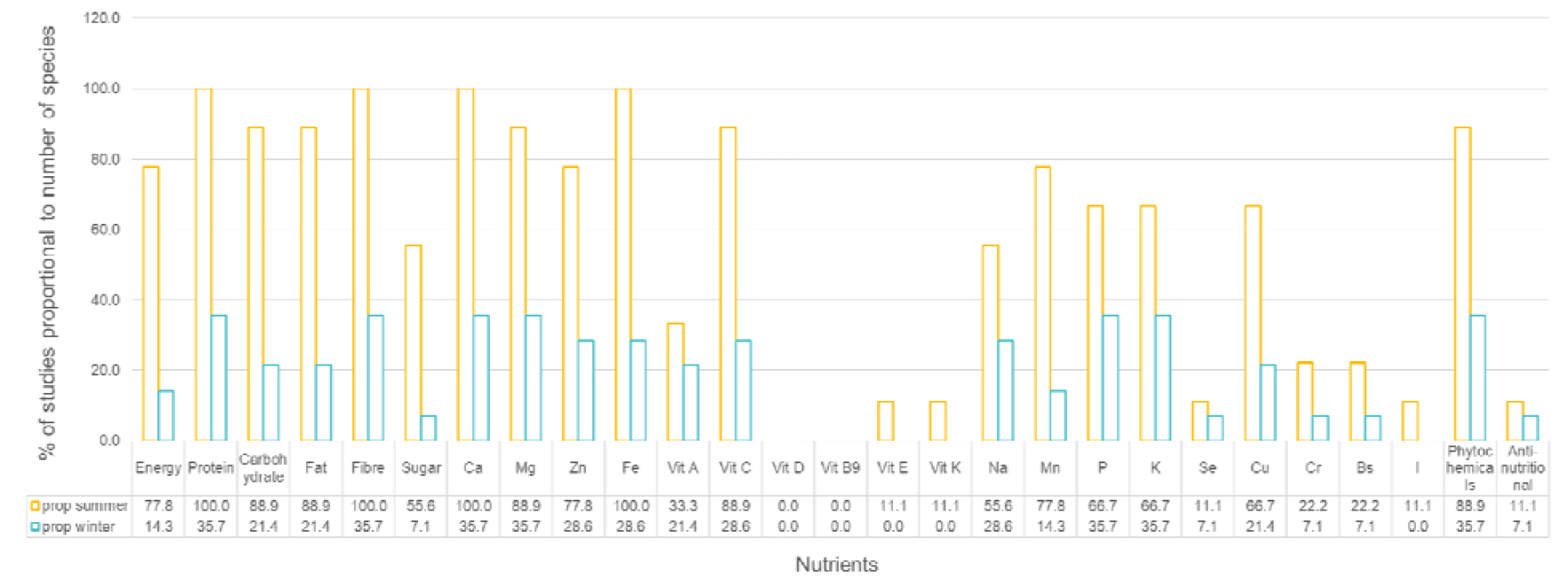
Research effort per nutrient relative to the number of species considered for that region (summer or winter rainfall) for which data existed. This is calculated as the percentage of incidences of a nutrient studied, proportional to the number of species considered for this region.

### B) Key Nutrients: nutritional values of key nutrients of those species studied

The results from the key nutrients analysis once again indicate that very little information exists on key nutrients for all species, with less data available for winter rainfall indigenous species. For summer rainfall species, *Mesembryanthemum cordifolium* has no data on any key nutrients whereas for winter rainfall species there are 7 species with no data on any key nutrients: *Prenia vanrensburgii, Trachyandra ciliata, Dasispermum suffruticosum, Searsia glauca, Mesembryanthemum guerichianum, Salvia aurea and Trachyandra divaricata*. The two exceptions amongst the winter rainfall species where more data exists on key nutrients include Carpobrotus edulis and *Mesembryanthemum crystallinum*.

Overall, a large range of key nutrient values is present. From the little data that is available, there is some indication that species indigenous to the GCFR may have the potential to contribute to the nutrition of diets (**Figure 4**). Examples include approximately up to ∼12% of the daily recommended intake for energy per portion, ∼8% protein, ∼80% fibre, ∼100% calcium, ∼80% Magnesium, and at least ∼25% Zinc, Iron and Vitamin C in only one portion (for details see **Supplementary Figures 1-2**). Specifically, *Mesembryanthemum crystallinum* also shows potential of contributing 10-45% of daily requirements of Vitamin C (27mg) per portion (80g) (**Supplementary Figure 2**). *Carpobrotus edulis* shows potential for contributing per 80g portion, an average of 6.8% energy, 7.1% protein, 55.8% fibre, 2.4% fat, 98.9% Calcium, 49.7% Magnesium, 23.2% Zinc, 21.4% Iron (**Supplementary Figures 1-2**).

**Figure 4.**
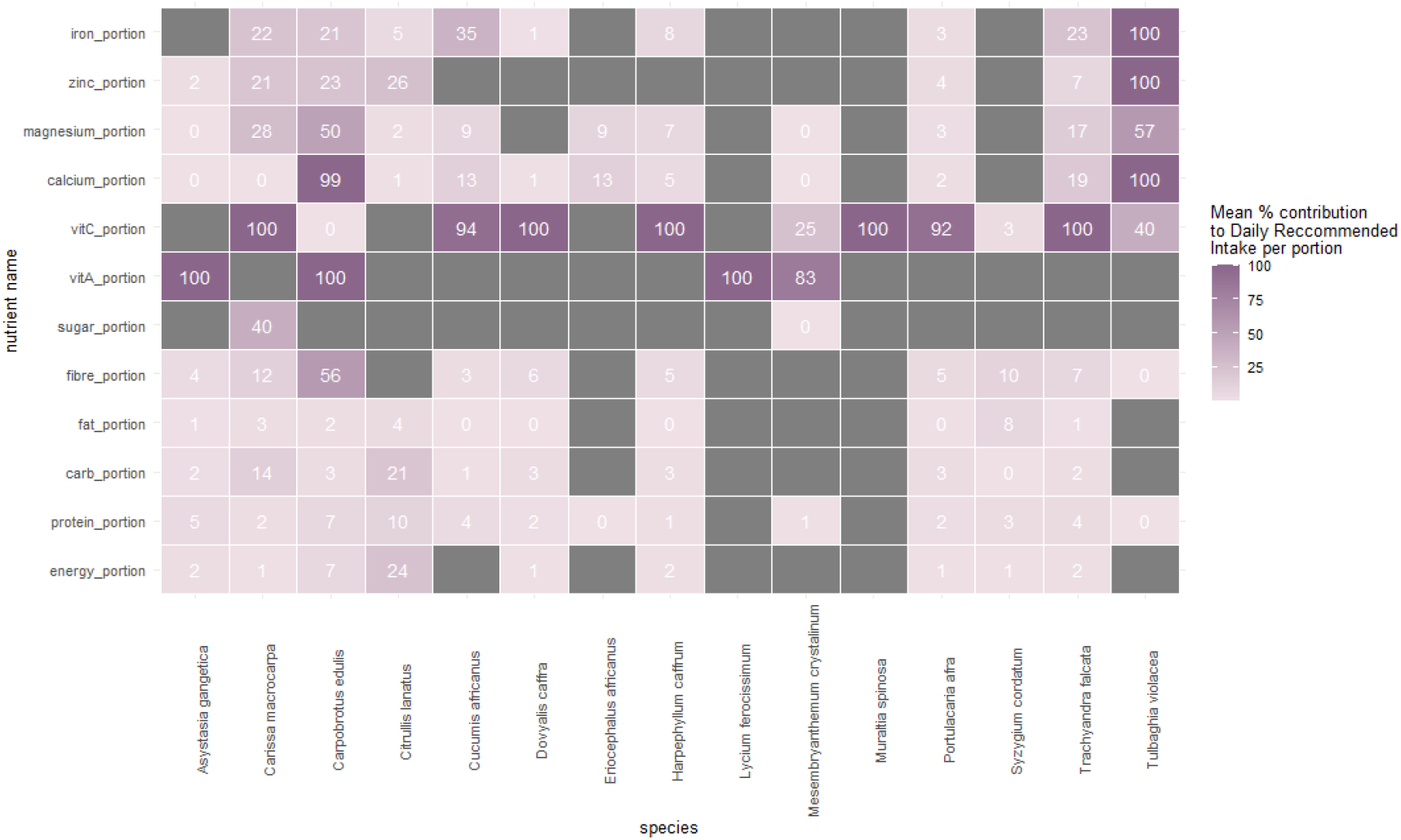
Heatmap indicating mean % contribution to Daily Recommended Intake of nutrients per portion for each species measured. Where this percentage was over 100%, we have capped this to 100% for illustrative purposes. See **supplementary Figures 1-2** to see full details.

Results indicate that generally, vitamins and minerals are high in the species for which studies exist, in particular *Carissa macrocarpa, Carpobrotus edulis, Trachyandra falcata and Tulbaghia violacea* (**Figure 4**). We also see that, despite few studies measuring Vitamin A, all species show high nutrient levels i.e. above 100% of daily recommended intake requirements per portion (also see **supplementary Figure 2**). Further, results show that some species are providing a higher mix of nutrients for example *C. macrocarpa, C. edulis, T. falcata, and T. violacea* **(Figure 5)**. Relatedly, **Figure 6** shows that *C. edulis* (a winter rainfall species) is the only species that provides all three nutrients (Iron, Zinc and Vitamin A) identified as deficient in the Western Cape.

**Figure 5.**
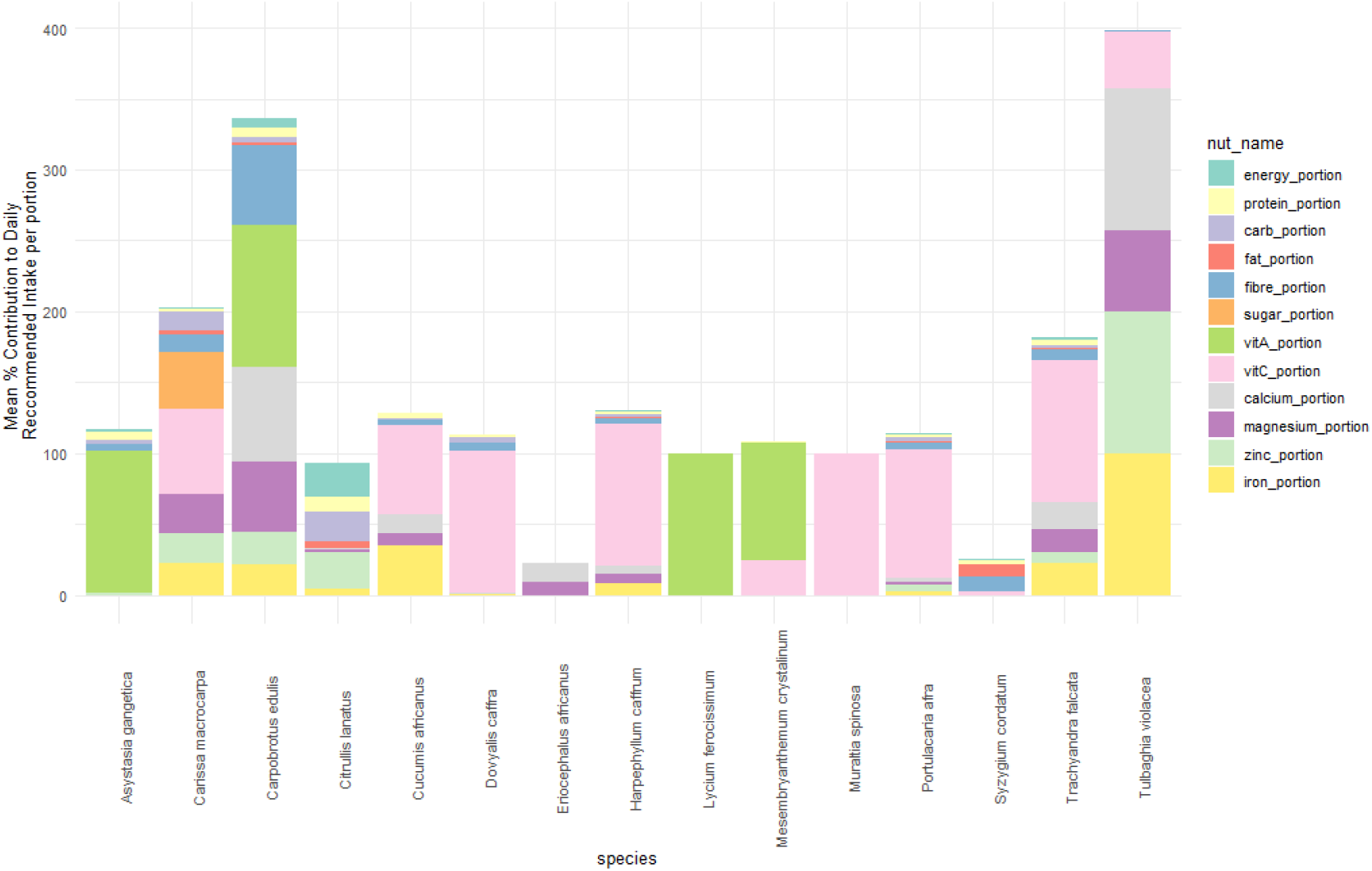
Mean nutrient contribution (%) to daily recommended intake per portion for species with nutritional data. Where this percentage was over 100%, we have capped this to 100% for illustrative purposes. See **supplementary figures 1-2** to see full details.

**Figure 6.**
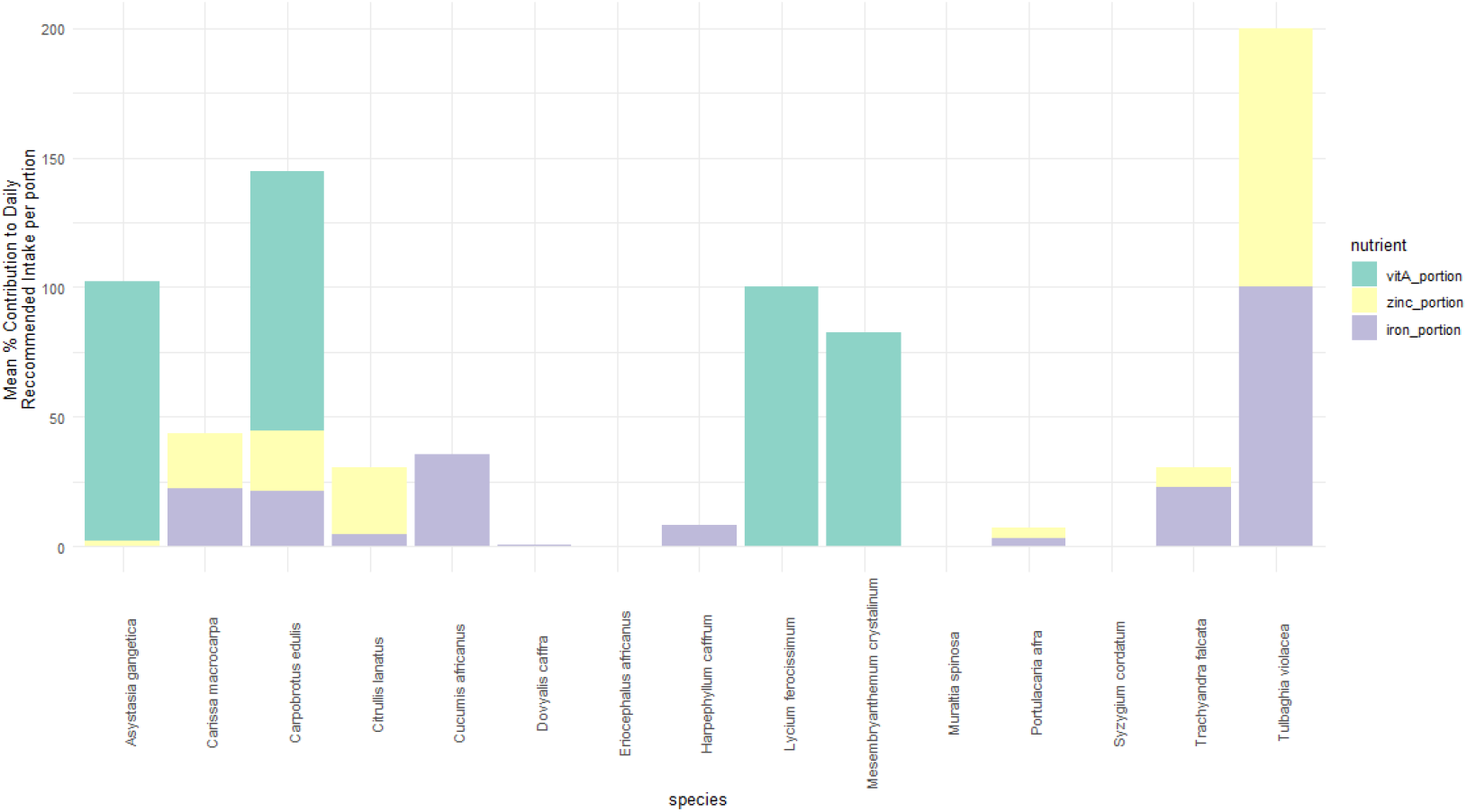
Mean nutrient contribution (%) to daily recommended intake per portion for species with nutritional data for a subset of nutrients (Vit A, Zinc, Iron) with known deficiencies in the Western Cape.

## 4. Discussion

### Research effort and key nutrients and priorities

Our study shows a lack of knowledge in nutritional status of local winter rainfall wild edibles except for 2 key species. This has likely contributed to greater momentum and motivation to cultivate summer rainfall species in the Western Cape, even if there exists great nutritional potential in more locally-adapted (water-wise, drought-tolerant and poor-soil-tolerant) species indigenous to the winter rainfall/Greater Cape Floristic Region. Our results therefore call for rapid upscaling of nutritional analysis on winter rainfall/GCFR species, as well as identifying priorities to establish baseline assessment of nutrition for those species with no nutritional data (e.g. *Prenia vanrensburgii, Tetragonia decumbens, Trachyandra cilliata, Dasispermum suffruticosum, Searsia glauca, Mesembryanthemum guerichianum, Salvia aurea* and *Trachyandra divaricata* species) and those nutrients with no data (Vitamin D and B9 for both regions, and Vitamin E, K and Iodine for the winter rainfall region).

For those species with existing data, *Carissa macrocarpa, Carpobrotus edulis, Trachyandra falcata*, and *Tulbaghia violacea* show both high levels of micronutrients (**Fig. 4**), and the greatest diversity of nutrients within a species (**Fig. 5**) and thus present species with potential to address micronutrient malnutrition through their consumption. *Tulbaghia violaceae* and *Trachyandra falcata* are particularly interesting to study further because despite having only a single study presenting data we see high contributions to nutrients: 1) Bvenura & Afolayan et al. (2017), find that *Tulbaghia violaceae* shows potential significant contributions per 10g portion consumed of Vitamin C (40%), calcium(>100%), magnesium(57%), zinc(>100%) and iron(>100%) and 2) Wehmeyer et al. (1986), find that *Trachyandra falcata* shows potential significant contributions per 80g portion consumed of protein (3.9%), fibre (7.5%), vitamin C (>100%), calcium (18.9%), magnesium (16.8%), zinc (7.4%) and iron (22.7%) in particular (**supplementary Figs. 1-2**). By extension, our results promote prioritisation of nutrient analysis for unstudied *Trachyandra ciliata* and *Trachyandra divaricata* due to relatedness to *Trachyandra falcata*, which likely have similar chemical properties. *Trachyandras* have also been identified as having gastronomical potential as they have been written about from a culinary perspective frequently in the past 5 decades (Coetzee & Miros, 2009, 2015; Leipoldt, 1976; Rood, 2008; Rusch, 2021, 2022; van der Merwe & de Villiers, 2015).

*Carpobrotus edulis* is one of the most studied species and shows great diversity and levels of nutrient contributions including for those nutrients of recognised deficiencies in the W. Cape (Faber M et al., 2011; Motadi et al., 2023), e.g. mean contributions to daily requirements of >100% Vitamin A, ∼23% Zinc and ∼21% Iron in just one portion (**Fig. 6**). This suggests a) the general promise of similar winter rainfall species and b) that this species specifically, could make a good candidate for a cultivation priority, provided we can determine its market potential, as we are already equipped with good baseline nutritional knowledge. A recent review (Winstead & Jacobson, 2024) identified *C. edulis* as a promising cultivation option due to its ability to thrive in arid, salty soils unsuitable for most agriculture. Additionally, both its leaves and flowers are edible, offering more than just the typically consumed fruit.

In terms of nutrient coverage per species as well as across species, our results identify some knowledge gaps. In the case of *Mesembryanthemum crystallinum*, there are only data for 5 out of 14 key nutrients. As this is a widely foraged and well-liked species (Rusch, 2021, 2022), a focus on other key nutrients including energy, carbohydrates, fibre, fat, zinc and iron, will be key for future studies on this species. *Lyceum ferocissimum* and *Muraltia spinosa* only had one nutrient studied each, both of which were over 100% contributions to daily requirements of Vitamin A and C respectively (**Fig. 4**), and thus present key species to investigate further nutritionally. Across species, Vitamin A warrants further investigation because contributions were mostly well over 100% contribution per portion for the four species studied (*A. gangetica, C. edulis, L. Ferocissum, M. crystallinum*). Furthermore, Vitamin D requires particular attention for the region, as it is a known seasonal deficiency (Middelkoop et al., 2022) but there was not a single study looking at this in species grown in the region.

### Why is it important we fill these gaps?

Species from this list show significant contributions to daily requirements of micronutrients such as calcium, magnesium, zinc, iron, vitamin C and A, of which Zinc, iron and Vitamin A were identified by Faber et al. (2011) as key deficiencies for the region. The feasibility of these species as potential solutions to nutrient deficiency will critically depend on consumer preference, their existing cultivation or the potential for easy household cultivation or sustainable harvesting. It is therefore key that we extend existing nutritional research for the edible plants grown in the Western Cape.

Addressing the knowledge gaps we identify in our study will also support the reintegration of these species into local foodways towards achieving economic, social, developmental and ecological goals.

Enhanced nutritional understanding of these species will ensure that it is possible to promote their uptake in homes by citing their health benefits in a targeted and informed manner. Greater market availability through food retailers (e.g. OZCF Market, Harvest of Hope, Living Soils) and restaurants will in turn stimulate income for the early adopter small scale farmers taking up growing these plants (Zhang & Dannenberg, 2022). Improved nutrition knowledge is also relevant to enabling policy to promote scaling the emerging innovation methods aimed at ensuring sustainable, regenerative and climate resilient cultivation. Socially and ecologically speaking, scaling the use of locally adaptive low resource use indigenous foods along with their understood nutrition benefits can contribute to reducing the negative impacts on people’s health and the region’s natural environments that mainstream crops are implicated in.

### Study limitations and future work

The absence of nutritional data identified in our study indicates a need for additional resources to further investigate the potential of local wild edible plants in addressing nutritional insecurity in the Western Cape. This will involve securing funding for targeted nutritional chemical analyses, but we should also explore alternative methods. For instance, innovative prediction methods for nutrients used by Cantwell-Jones et al. (2022) for Vitamin B prediction using phylogenies could be an initial step in pinpointing which species warrant further analysis. In fact, two priority species identified by our study, *Trachyandra falcata* and *Tulbaghia violacea*, were predicted by Cantwell-Jones et al. (2022) to have high levels of Vitamin B, a nutrient commonly deficient but key for a healthy nervous system.

A further limitation that requires some attention is the standardisation of nutrient data reported in these studies. Protocols and units varied per study making it difficult to compare certain studies and many had to be excluded as a result of poor reporting/lacking units etc. Our study therefore calls for a standardisation in protocol and reporting for nutritional analysis to advance this field of study and thus the impact this research can have for reintegrating these species into food systems as nutrition security solutions.

Due to the region’s incredible biodiversity, there are many more potential candidates to assess for their nutritional value that were not included in this study. For example, there are ∼300 species of *Searsia* that occur across South Africa; of which some have been cited as used historically in ethnobotany records (De Vynck et al., 2016) and found occurring at Archaeological sites of habitation (van Wijk et al., 2017) - thus indicating that these species are likely to be nutritious. They are related to Mediterranean sumac which is a widely used spice. Similarly there are multiple species of *Tetragonia* widespread across winter and summer rainfall regions and found in ethnobotanical records (van Wijk et al., 2017), which are not only recognized as edible by rural communities but are also favoured by animals, suggesting potential use as animal fodder (Personal observations by author L. Rusch). As identified in our study *Trachyandra* species beyond *T. falcata*, warrant further investigation due to the gastronomic familiarity of these species being prepared and eaten in a similar manner to asparagus or long-stem broccoli (Rusch, 2022) as well as early evidence of ease of cultivation (Rusch, 2021).

A key element required to reintegrate the species we identify in this study as having nutritional potential and which is still largely lacking, is a comprehensive understanding of the market potential or consumer preference or value of species. Recent research (Zhang & Dannenberg, 2022) identified the realised challenges for the marketisation of these species including limited seed/cutting access for growers, limited capacity of growers, competition with subsidised conventional production and limited distribution options. However, this study also highlighted the great potential for income generation and environmental adaptation which these species present.

This links to the final consideration that will be required to upscale this work and ensure its long-term sustainability: to understand the climate change resilience of these species. It is already expected that these species may outperform conventional crops due to inherent adaptations to drought, poor soils and fires which characterise the region. This will become increasingly important under predicted increased intensity of droughts and fires and overall aridification of the Western Cape under future climate change. It will be important to quantify this resilience per species through multi-pronged approach of understanding future suitability through species distribution modelling as well as through cultivation trials under varying conditions simulating future conditions. Currently, there’s a lack of understanding about the climate change resilience of most edible species in the region. However, there are promising signs from both modeling, which predicts an increase in species richness of wild food plants despite habitat suitability declines, and cultivation studies, such as *M. crystallinum* and *T. decumbens*, which show resilience to water stress (Tembo-Phiri et al., 2019).

Through this process of quantifying the potential of species as future food solutions, caution must be taken when highlighting their potential to avoid promotion of unsustainable harvesting through foraging. Our recommendations are therefore strictly aimed through a lens of upscaling subsistence growing and establishing cultivation for marketisation in order to protect species populations in the wild, rather than exacerbate conservation concerns for the region by encouraging foraging.

## 5. Conclusions

There are a number of species grown in the Western Cape that are key for further nutritional analysis due to their potential nutrient contributions despite zero or limited assessments of nutrition currently in existence. Those species indigenous to the winter rainfall/ Greater Cape Floristic region are particularly under researched despite evidence suggesting their high nutritional potential (e.g. *Trachyandra falcata*). We identify *Carpobrotus edulis* as a key indigenous species to most imminently prioritise for future cultivation and marketisation because there is comprehensive knowledge of its nutritional status which shows that it has high nutrition contributions to make.

However, it is likely a combination of species of those grown in the GCFR, including the understudied indigenous species that will provide key resources for addressing nutritional security of the Western Cape. We have demonstrated that within these wild edibles there are opportunities to address recognised nutritional deficits in the region (e.g. *C. edulis* and *T. violacea* providing significant contributions of vitamin A, Zinc and Iron). It is thus critical to expand this research to determine opportunities for marketisation and cultivation, recognising that highlighting species should not encourage new foraging so that we can sustainably use and preserve the regions unique edible biodiversity.

## Author contributions

LP, LR and NK conceived the study, MR, NK, FS and EvdM collected the data. NK analysed the data. NK drafted the manuscript. NK, LP and LR edited the final manuscript.

## Acknowledgements

The authors would like to thank Dr. Rhoda Malgas for helpful discussions about this research.

## Data availability and disclosure statement

Data used to generate this analysis are available from the authors upon reasonable request. The authors report there are no competing interests to declare.

## Supplementary Material

Separate text file.

## References

Akinola, R., Pereira, L. M., Mabhaudhi, T., de Bruin, F. M., & Rusch, L. (2020). A review of indigenous food crops in Africa and the implications for more sustainable and healthy food systems. In Sustainability (Switzerland) (Vol. 12, Issue 8). MDPI. 10.3390/SU12083493

Borelli, T., Hunter, D., Powell, B., Ulian, T., Mattana, E., Termote, C., Pawera, L., Beltrame, D., Penafiel, D., Tan, A., Taylor, M., & Engels, J. (2020). Born to eat wild: An integrated conservation approach to secure wild food plants for food security and nutrition. In Plants (Vol. 9, Issue 10, pp. 1–37). MDPI AG. 10.3390/plants9101299

Botha, M. S., Cowling, R. M., Esler, K. J., de Vynck, J. C., Cleghorn, N. E., & Potts, A. J. (2020). Return rates from plant foraging on the Cape south coast: Understanding early human economies. Quaternary Science Reviews, 235. 10.1016/j.quascirev.2019.106129

Bvenura, C., & Afolayan, A. J. (2017). Tackling food and nutrition insecurity using leafy wild vegetables: The nutritional compositions of some selected species. In A. Méndez-Vilas (Ed.), Science within Food: Up-to-date Advances on Research and Educational Ideas (p. 243). FORMATEX.

Cantwell-Jones, A., Ball, J., Collar, D., Diazgranados, M., Douglas, R., Forest, F., Hawkins, J., Howes, M. J. R., Ulian, T., Vaitla, B., & Pironon, S. (2022). Global plant diversity as a reservoir of micronutrients for humanity. Nature Plants, 8(3), 225–232. 10.1038/s41477-022-01100-6

Coetzee, R., & Miros, V. (2009). Koekemakranka. Lapa Uitgewers.

Coetzee, R., & Miros, V. (2015). A Feast From Nature. Penstock Publishing.

De Vynck, J. C., Van Wyk, B. E., & Cowling, R. M. (2016). Indigenous edible plant use by contemporary Khoe-San descendants of South Africa’s Cape South Coast. South African Journal of Botany, 102, 60–69. 10.1016/j.sajb.2015.09.002

DEA (Department of Environmental Affairs). (2013). Long Term Adaptation Scenarios Flagship Research Programme (LTAS) for South Africa. Climate Change Implications for the Biodiversity Sector in South Africa. Department of Envionmental Affairs., South Africa.

Faber M S. J.,, Witten, C., & Drimie, S. (2011). Review: Community-based agricultural interventions in the context of food and nutrition security Community-based agricultural interventions in the context of food and nutrition security in South Africa. South African Journal of Clinical Nutrition, 24(1).

Gold, K., & McBurney, R. P. H. (2012). Conservation of plant diversity for sustainable diets. In B. Burlingame & S. Dernini (Eds.), Sustainable diets and biodiversity: Directions and solutions for policy, research and action (pp. 108–115). Food and Agriculture Organization of the United Nations; Bioversity International.

Leipoldt, C. L. (1976). Leipoldt’s Cape Cookery. W.J. Flesch Publications.

Middelkoop, K., Walker, N., Stewart, J., Delport, C., Jolliffe, D. A., Nuttall, J., Coussens, A. K., Naude, C. E., Tang, J. C. Y., Fraser, W. D., Wilkinson, R. J., Bekker, L. G., & Martineau, A. R. (2022). Prevalence and Determinants of Vitamin D Deficiency in 1825 Cape Town Primary Schoolchildren: A Cross-Sectional Study. Nutrients, 14(6). 10.3390/nu14061263

Motadi, S. A., Zuma, M. K., Freeland-Graves, J. H., & Mbhenyane, G. X. (2023). Iron and zinc status of children aged 3 to 5 years attending Early Childhood Development centres in Venda, South Africa. Ecology of Food and Nutrition. 10.1080/03670244.2023.2210502

Motti, R. (2022). Wild Edible Plants: A Challenge for Future Diet and Health. In Plants (Vol. 11, Issue 3). MDPI. 10.3390/plants11030344

Pironon, S., Ondo, I., Diazgranados, M., Allkin, R., Baquero, A. C., Cámara-Leret, R., Canteiro, C., Dennehy-Carr, Z., Govaerts, R., Hargreaves, S., Hudson, A. J., Lemmens, R., Milliken, W., Nesbitt, M., Patmore, K., Schmelzer, G., Turner, R. M., van Andel, T. R., Ulian, T., … Willis, K. J. (2024). The global distribution of plants used by humans. https://www.science.org

Rood, B. (2008). Kos uit die veldkombuis. Protea Boekhuis.

Rouget, M., Barnett, M., Cowling, R. M., Cumming, T., Daniels, F., Hoffman, M. T., & Rebelo, T. (2014). Chapter 14: Conserving the Cape Floristic Region. In N. Allsopp, J. F. Colville, & A. Verboom (Eds.), Fynbos: Ecology, Evolution, and Conservation of a Megadiverse Region (p. 321). Oxford University Press.

Rusch, L. (2021). Cape Wild Foods: a Growers Guide (2nd ed.). The Sustainability Institute.

Rusch, L. (2022). Cape Wild Foods: a Cooks Guide. The Sustainability Institute.

Singels, E., Parkington, J., & Esler, K. J. (2020). The role of geophytes in Stone Age hunter-gatherer subsistence and human evolution in the Greater Cape Floristic Region. University of Cape Town.

Singels, E., Potts, A. J., Cowling, R. M., Marean, C. W., De Vynck, J., & Esler, K. J. (2016). Foraging potential of underground storage organ plants in the southern Cape, South Africa. Journal of Human Evolution, 101, 79–89. 10.1016/j.jhevol.2016.09.008

Snijman, D. (2013). Plants of the Greater Cape Floristic Region. 2: The Extra Cape flora. (D. Snijman, Ed.; Vol. 2). South African National Biodiversity Institute.

Solomons, N., Kruger, H. S., & Pouane, T. R. (2019). Addressing non-communicable diseases in the Western Cape, South Africa. In Journal of Public Health Research (Vol. 8).

Tembo-Phiri, C., Phiri, E. E., & Rusch, L. C. (2019). Edible Fynbos Plants: A Soil Types and Irrigation Regime Investigation on Tetragonia decumbens and Mesembryanthemum crystallinum [MSc, University of Stellenbosch]. https://scholar.sun.ac.za

UNICEF/WHO/World Bank. (2022). Global Nutrition Report. Https://Globalnutritionreport.Org/Resources/Nutrition-Profiles/Africa/Southern-Africa/South-Africa/.

Uusiku, N. P., Oelofse, A., Duodu, K. G., Bester, M. J., & Faber, M. (2010). Nutritional value of leafy vegetables of sub-Saharan Africa and their potential contribution to human health: A review. In Journal of Food Composition and Analysis (Vol. 23, Issue 6, pp. 499–509). 10.1016/j.jfca.2010.05.002

van der Merwe, K., & de Villiers, J. (2015). Strandveldfood: A West Coast Odyssey. Sunbird Publishers.

van Wijk, Y., Tusenius, M. L., Rust, R., Cowling, R. M., & Wurz, S. (2017). Modern vegetation at the klasies river archaeological sites, tsitsikamma coast, South-Eastern Cape, South Africa: A reference collection. Plant Ecology and Evolution, 150(1), 13–34. 10.5091/plecevo.2017.1286

Wehmeyer, A. (1986). Edible wild plants of Southern AfricafJ: Data on the nutrient contents of over 300 species.

Welcome, A. K., & Van Wyk, B. E. (2019). An inventory and analysis of the food plants of southern Africa. South African Journal of Botany, 122, 136–179. 10.1016/j.sajb.2018.11.003

Winstead, D. J., & Jacobson, M. G. (2024). Storable, neglected, and underutilized species of Southern Africa for greater agricultural resilience. In Plant-Environment Interactions (Vol. 5, Issue 4). John Wiley and Sons Inc. 10.1002/pei3.70004

Zhang, M., & Dannenberg, P. (2022). Opportunities and Challenges of Indigenous Food Plant Farmers in Integrating into Agri-Food Value Chains in Cape Town. Land, 11(12). 10.3390/land11122267

Ziervogel, G., New, M., Archer van Garderen, E., Midgley, G., Taylor, A., Hamann, R., Stuart-Hill, S., Myers, J., & Warburton, M. (2014). Climate change impacts and adaptation in South Africa. Wiley Interdisciplinary Reviews: Climate Change, 5(5), 605–620. 10.1002/wcc.295

