## Supplementary tables and figures for "Harnessing the Potential of Cape Wild Edible Plants: Nutritional insights, Gaps and Priorities"

**Supplementary Table 1**: Species list of species featured in this study with full list of common names and family.

| **Family name** | ***Scientific name*** | **Common name(s)** |
| --- | --- | --- |
|  | **LEAFY GREENS ,SUCCULENTS AND VEGETABLES** | |
| ***Aizoaceae*** | *Tetragonia decumbens* | Dune spinach, duinespinasie |
|  | *Mesembryanthemum crystallinum* | Soutslaai, Ice Plant |
|  | *Mesembryanthemum guerichianum* | Brakslaai |
|  | *Mesembryanthemum cordifolium* | Sun Rose |
|  | *Prenia vanrensburgii* | See pampoen, cape spinach, strandspinasie |
| **Portulacacea** | *Portulacaria afra* | Spekboom, porkbush, igwanitsha (isiXhosa), isicococo, isidibiti esikhulu (isiZulu) |
| **Acanthaceae** | *Asystasia gangetica* | Forest Foxglove, tropical primrose, Chinese Violet |
| **Asphodeloideae** | *Trachyandra ciliata* | Veldkool; wilde blomkool; moretete, mototo, (Sotho) |
|  | *Trachyandra falcata* | Veldkool |
|  | *Trachyandra divaricata* | Sandkool |
| **Cucurbitaceae** | *Cucumis africanus* | wild cucumber, springbok cucumber, Monyaku (Pedi/ Sotho), wilde agurkie, tshinyagu (venda) doringkomkommertjie, mthangazana/ Ithangazana (Xhosa) isende-lenja or uselwa-lwemamba in Swazi and Zulu; bitter apple; bitterappel |
| **Cucurbitaceae** | *Citrullis lanatus subsp lanatus var citroides* | tsamma; makataan |
|  | **HERBACIOUS CULINARY WILD HERBS (NO WOODY STEM)** | |
| **Apiaceae** | *Dasispermum suffruticosum* | Dune Celery; duineselderei |
| **Alliaceae** | *Tulbaghia violaceae* | society garlic, wild garlic |
|  | **SHRUB LIKE CULINARY WILD HERBS (DEVELOP WOODY STEM)** | |
| **Asteraceae** | *Eriocephalus africanus* | Wild rosemary; Cape snowbush; kapokbos; |
| **Lamiaceae** | *Salvia aurea* | Brown sage |
|  | **SOFT FRUITS (CONTAIN SEVERAL SEEDS)** | |
| **Solanaceae** | *Lycium ferocissimum* | Slangbessie; African boxthorn |
| **Apocynaceae** | *Carissa macrocarpa* | Num num; Amathungulu (fruit in isiZulu); Natal plum; umthungulu (isiZulu, isiXhosa) |
| **Saliaceae** | *Dovyalis caffra* | Kei apple; wild apricot; appelkoosdoring; wilde-appelkoos;umqokolo (isiXhosa; isiZulu); umqakalo (isiZulu); amaqogolo, umkokolo; umquokolo (Ndebele); mahlono (Pedi); mukokolo; munhunguru; mutumotote (Shona); mohloni (Sotho) |
|  | **DRUPES (FLESH AROUND SINGLE HARD SEED)** | |
| **Anacardiaceae** | *Searsia glauca* | blue kunibush; bloukoeniebos |
| **Polygalaceae** | *Muraltia spinosa* | skilpadbessie; tortoise berry; cargoe |
| **Myrtaceae** | *Syzygium cordatum* | Waterberry; waterbessie; umdoni (isiZulu, isiXhosa), umswi (isiXhosa); umdoni, umswi (Ndebele); , umngcosi (isiSwati); mukute, muwototo (Shona); muhlu, munonyanansi (Tsonga); timuhlu (Shangaan) |
| **Anacardiaceae** | *Harpephyllum caffrum* | Wild Plum; umgwenya (isiXhosa, isiZulu, Swati); wildepruim (Afrikaans); mmedibibi (Pedi) |

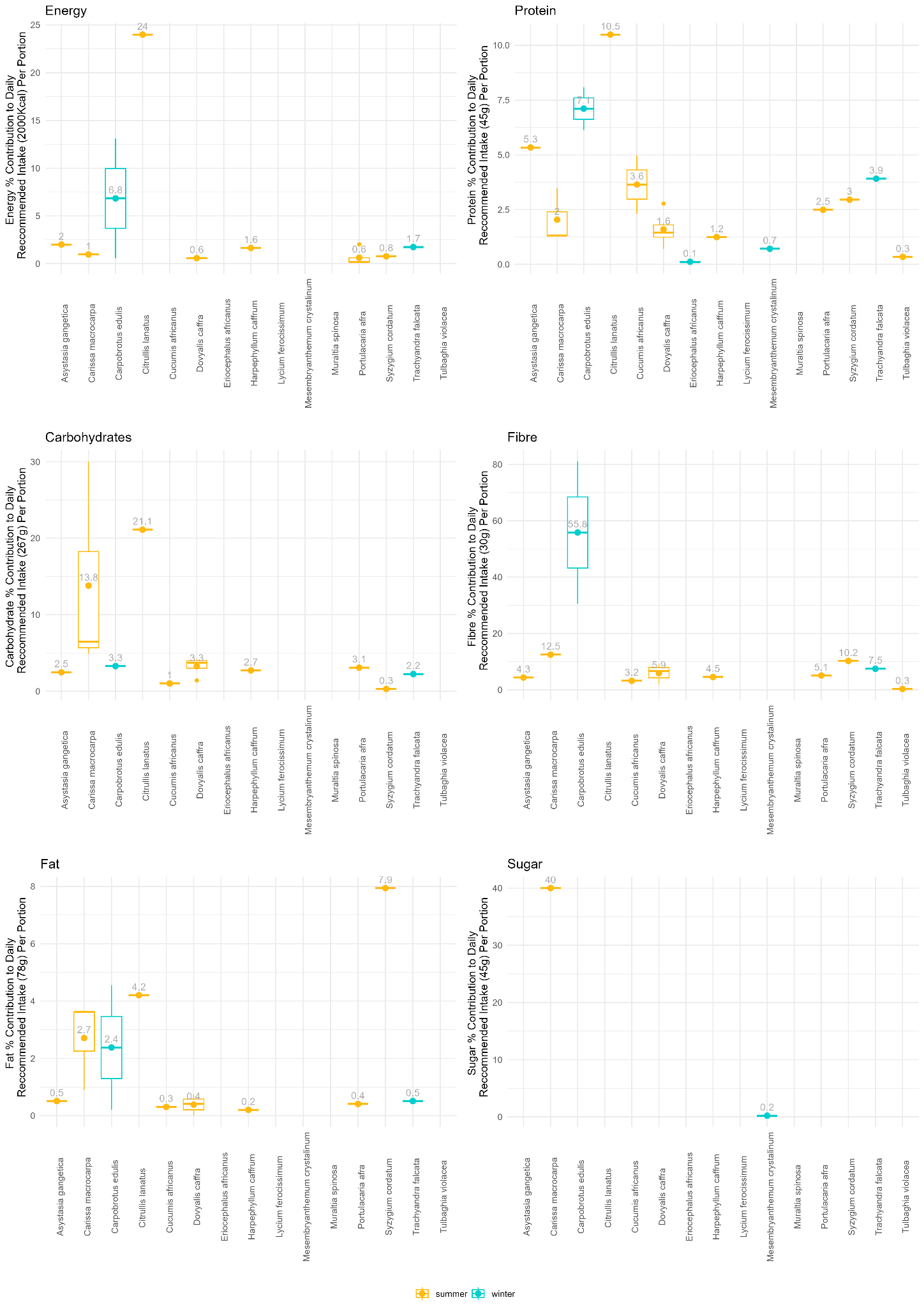

***Supplementary Figure 1:*** *Percentage contribution by species of each macronutrient to the daily recommended intake of that nutrient per portion. Portion is unique per species (calculated as an average portion of either vegetable-like/succulent leaves (80g), fruit/drupe/berry (80g) or herbs/aromatics (10g).*

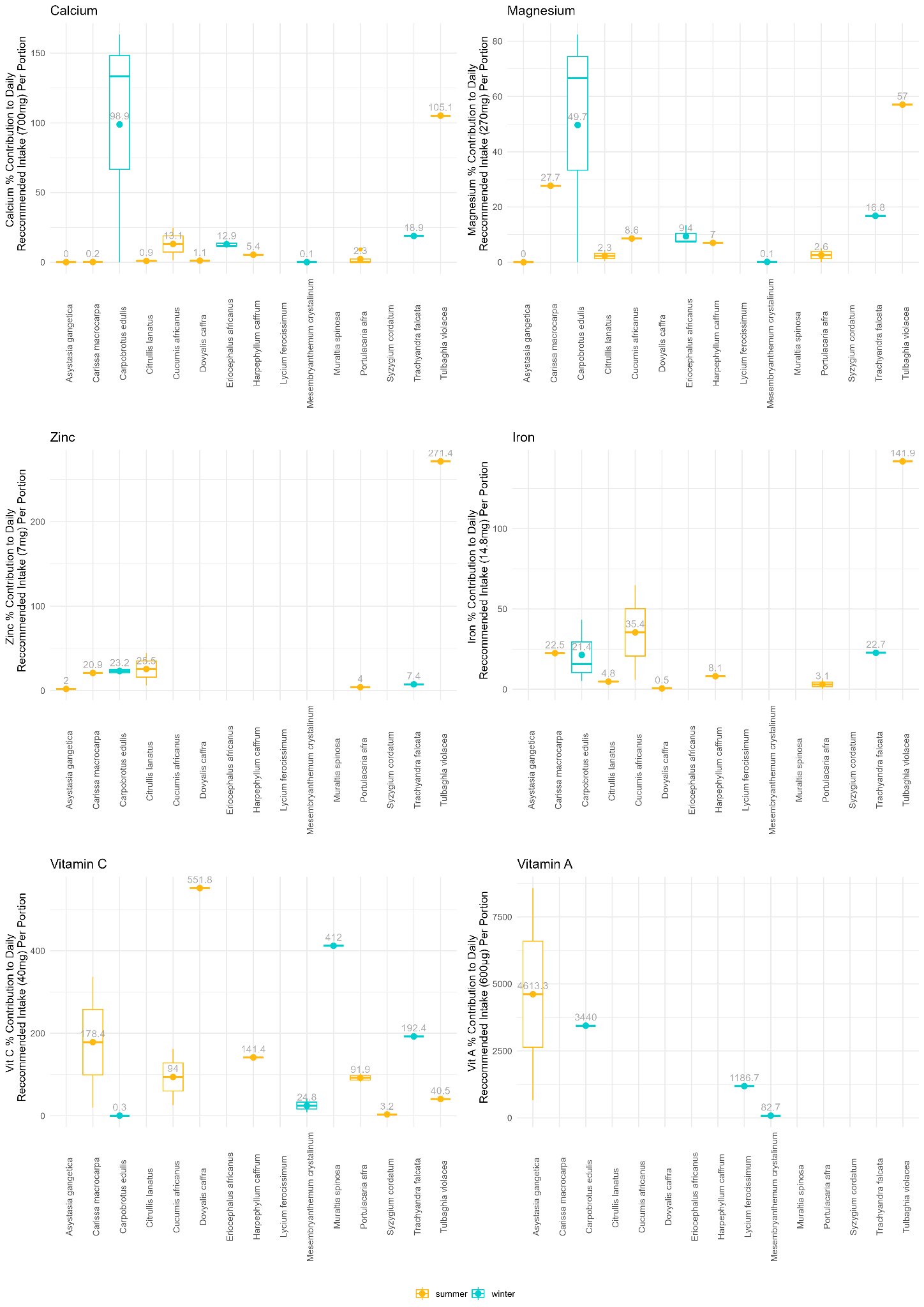

***Supplementary Figure 2:*** *Percentage contribution by species of each micronutrient to the daily recommended intake of that nutrient per portion. Portion is unique per species (calculated as an average portion of either vegetable-like/succulent leaves (80g), fruit/drupe/berry (80g) or herbs/aromatics (10g).*
